# Messenger-RNA Modification Standards and Machine Learning Models Facilitate Absolute Site-Specific Pseudouridine Quantification

**DOI:** 10.1101/2022.05.06.490948

**Authors:** Amr Makhamreh, Sepideh Tavakoli, Howard Gamper, Mohammad Nabizadehmashhadtoroghi, Ali Fallahi, Ya-Ming Hou, Sara H. Rouhanifard, Meni Wanunu

## Abstract

Enzyme-mediated chemical modifications to mRNA are important for fine-tuning gene expression, but they are challenging to quantify due to low copy number and limited tools for accurate detection. Existing studies have typically focused on the identification and impact of adenine modifications on mRNA (m^6^A and inosine) due to the availability of analytical methods. The pseudouridine (Ψ) mRNA modification is also highly abundant but difficult to detect and quantify because there is no available antibody, it is mass silent, and maintains canonical basepairing with adenine. Nanopores may be used to directly identify Ψ sites in RNAs using a systematically miscalled base, however, this approach is not quantitative and highly sequence dependent. In this work, we apply supervised machine learning models that are trained on sequence-specific, synthetic controls to endogenous transcriptome data and achieve the first quantitative Ψ occupancy measurement in human mRNAs. Our supervised machine learning models reveal that for every site studied, different signal parameters are required to maximize Ψ classification accuracy. We show that applying our model is critical for quantification, especially in low-abundance mRNAs. Our engine can be used to profile Ψ-occupancy across cell types and cell states, thus providing critical insights about physiological relevance of Ψ modification to mRNAs.

## Introduction

RNA modifications are critical for cellular function, as demonstrated by their requirement for proper folding and stability of tRNA and rRNA where they were first discovered^1^. By analyzing these highly expressed RNAs, over 100 different types of RNA modifications have been discovered and characterized using analytical tools such as mass spectrometry^2^ and thin-layer chromatography^3^. As sequencing technologies have developed, many of these modifications have also been identified on messenger RNAs such as inosine^4^, N6-methyladenine (m^6^A)^5,6^ and pseudouridine^7,8^. Next-generation sequencing studies have begun to unravel the role mRNA modifications play in finetuning gene expression. However, identifying the precise modification site within the mRNA sequence and the fractional occupancy (i.e., fraction of copies with that modification) is a daunting task^9^. Low mass abundance of individual mRNA species in transcriptomes precludes the use of existing methods such as mass spectrometry, and chemical labeling methods are not quantitative. Pseudouridine (Ψ) is among the most highly represented mRNA modifications and is typically detected using biochemical labeling methods. Pseudouridine-modified mRNAs are more resistant to RNAsemediated degradation^10^ and also have the potential to modulate immunogenicity^11^ and enhance translation^12^ *in vivo*. During the COVID-19 pandemic, Ψ has taken the spotlight due to the inclusion of the methylated Ψ analog, N1-methylpseudouridine, in the Moderna^13^ and Pfizer^14^ mRNA vaccines for SARS CoV-2.

Tools for high-confidence, transcriptome-wide identification of RNA modifications, in particular Ψ, have been somewhat limited due to a lack of chemical specificity and proper ‘gold-standard’ controls for accurate benchmarking. Coupling next generation sequencing (NGS) with modification-specific chemicals (i.e. CMC^7,8,15^ or bisulfite sequencing^16^) can be used to identify sites, but due to a reliance on cDNA amplification this method is not quantitative and prone to bias. Thus, there is little overlap between the identified sites using each method. Moreover, since these methods rely on base deletion or read termination for detection, tandem modifications on the same transcript cannot be detected. To this end, non-destructive detection of native RNA molecules is the most attractive approach for reading epitranscriptome landscapes. The most promising method thus far has been direct RNA nanopore sequencing, which offers the ability to preserve full-length RNA structural information^17^. In this method, an RNA strand is ratcheted through a nanopore and the ion current signal produces reports on its sequence by sequentially reading a string of k-mers (k=5). Variance in the signals from the consensus expected signals of unmodified bases can be used to identify modifications. We and others have recently shown that these signal anomalies produce systematic base-calling errors at or near the site of Ψ modification^18–22^. In addition, prediction models have been developed to improve modification calls by leveraging features like deviations in the expected ionic current, systematic base mismatches, changes in base quality score, and insertion/deletion rates^19,20,23^.

Previous works have reinforced the confidence of Ψ-site calling from direct RNA sequencing data, which often presents itself as a U-to-C mismatch error. However, the training data sets contain satellite modifications close to the Ψ-site, which can introduce undesirable noise and reduced accuracies when training a 5-mer specific model. For example, *in vitro* transcribed RNA constructs bearing all combinations of Ψ-containing 5-mers have been used to generate nanopore-based training data for Ψ modifications^24^. While cost-effective, since all U sites have been replaced with Ψ, training a Ψ detection model for regions in native RNAs where the 5-mer sequence contains more than one U site is not feasible. Recent work by Fleming et al.^21^ involved the design of synthetic constructs that separate Ψ-sites by ∼25 nucleotide spacers to remove the effects of satellite modifications at the protein motor and pore, allowing them to test signal dwell time corresponding to when Ψ is located in the helicase motor as another feature for discrimination. With these constructs, U-to-C mismatch rates varied from 10% to 97% across 15 different Ψ-modified 5-mers. However, these constructs also lack 5-mers with canonical U’s adjacent to Ψ, which are often found in the transcriptome.

To address these challenges and improve the accuracy of Ψ detection by direct RNA sequencing, we performed a meta-analysis of four synthetic constructs bearing a singly- modified Ψ within an endogenous mRNA sequence. These four sites were flagged by our Ψ-detection algorithm^22^. Interestingly, we found that the U-to-C mismatch rates for the 100% Ψ-modified constructs varied from 30% to 70%, and further, that these depend on the specific k-mer and sequence context. If mismatch errors were fully quantitative, we would expect to see 100% U-to-C mismatch in all constructs; however, the method is highly sequence-specific, and is therefore only effective at identifying modification sites^19^.

We were interested to see whether our synthetic constructs can be used for training 5-mer specific models that can accurately quantify Ψ occupancy at identified sites in native mRNA. Toward this, we developed and tested a computational tool that can train supervised-machine learning (ML) models on nanopore-based features derived from our four synthetic Ψ-modified constructs to subsequently quantify Ψ occupancy at these specific locations in native HeLa mRNA transcripts. We find that Ψ discrimination with 5-mer-specific ML models trained with basecalling and raw signal features prepared from labeled 100% and 0% Ψ-modified synthetic reads can achieve accuracies above 90%, even at low Ψ occupancies. In addition, we found that the combination of features conducive for classification accuracy depends on the sequence context of the Ψ-modified 5-mer region. Finally, we applied these trained models and achieved the first demonstration of site-specific Ψ quantification in human mRNAs.

## Results

### Supervised Machine Learning on Ψ-modified Synthetic Transcripts

Our pipeline for quantitative Ψ profiling is shown in Figure 1. We recently developed a set of four synthetic RNA control standards that bear established and putative Ψ-modification positions in the HeLa transcriptome^22^. Briefly, two of the constructs, *MCM5 (chr22: 35424407,UGUAG)* and *PSMB2 (chr1: 35603333, GUUCG)*, have been validated by CeU-seq^15^, Ψ-seq^8^, and RBS-seq^16^, while the other two, *MRPS14 (chr1: 175014468, ACUUA), PRPSAP1 (chr17: 76311411, GAUUG)* were indirectly detected *de novo* by observing a significantly high U-to-C mismatch error in direct RNA nanopore sequencing. We will subsequently refer to each of these constructs by the gene name and omit the modification position. Briefly, 100% Ψ-modified (syn-Ψ) standards bearing a Ψ-modification were generated (**Fig. 1a**, yellow box), as well as the corresponding, sequence-matched, unmodified transcript (syn-U). We were interested to compare different supervised machine learning (ML) models and find the optimal model that can accurately and quantitatively classify Ψ-sites in the synthetic controls. To determine the most optimal combination of features we extracted basecalling and raw signal features at both local (Ψ-site) and remote (upstream) for a total of 60 features. Understanding which signal features optimize Ψ classification in mRNA is a crucial step when developing a quantification method. Thus, we extracted 60 signal features from each synthetic read that passed the 2^nd^ filtration stage (see methods) in both the syn-U library and syn-Ψ library (**Fig. 1c**) using *nanopolish*^25^. These features were subsequently used to generate and test different supervised machine learning classifiers. The features were basecalls, which included deletions, quality scores of positions -2, -1, 0 [U/Ψ], +1, +2, current mean, current standard deviation, dwell time, and Fourier coefficients 2 and 3 (FC2 and FC3) of the 5-mers where Ψ is positioned at - 2, -1, 0, +1, +2, and going 12 bases upstream (3’ direction) to the 5-mers when Ψ is at the protein motor (−14, -13, -12, -11, -10) we also extracted their current mean, current standard deviation, dwell time, and FC2, FC3. Raw signal features were extracted and compiled into one dataframe from Fast5 files using the *eventalign resquiggle* tool from nanopolish^25^ (**Supplementary Table S5**).

**Figure 1.**
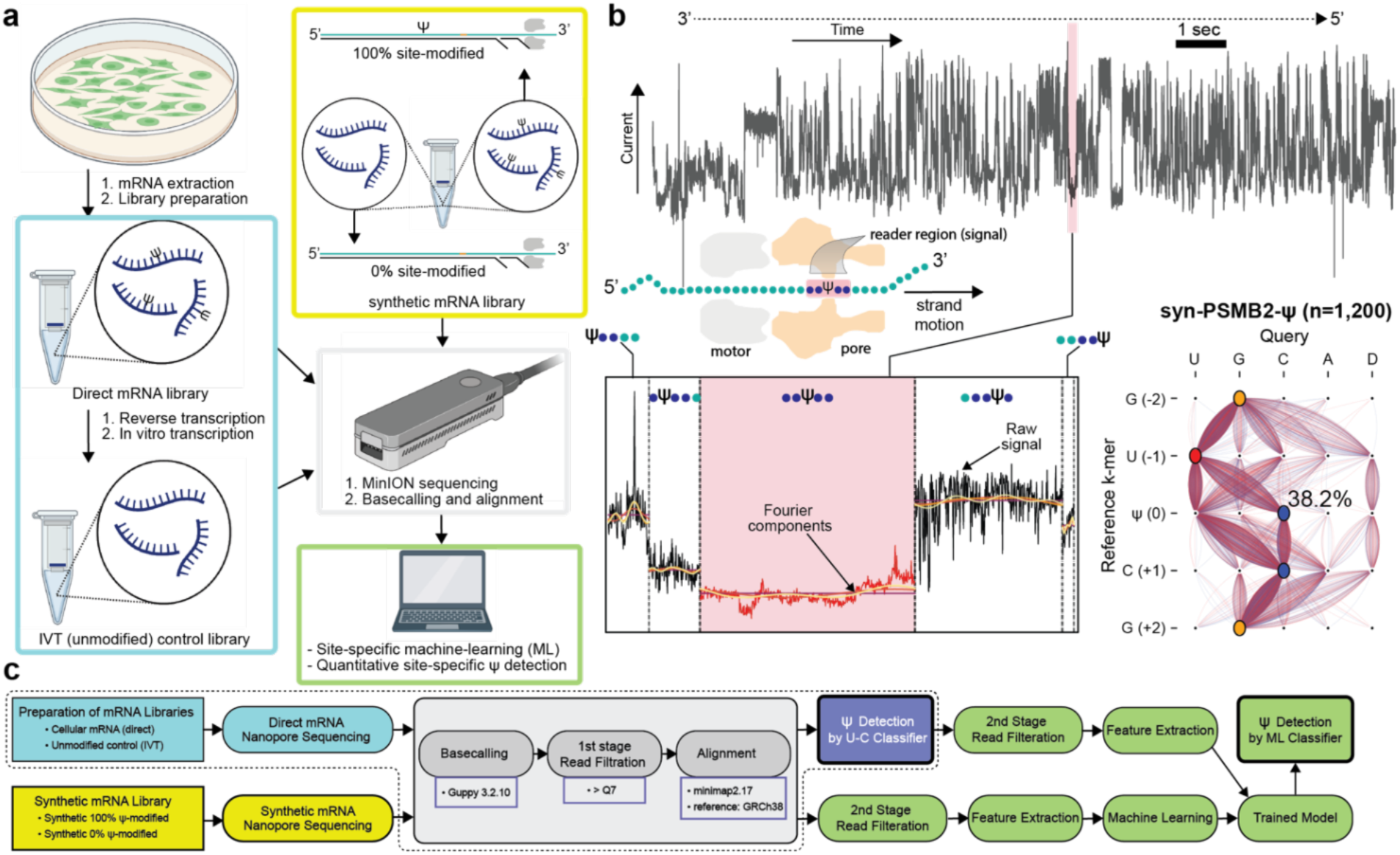
Synthetic RNA pipeline for quantitative pseudouridine profiling. a, A typical RNA processing pipeline from cells (left) or a synthetically prepared library (right). After RNA extraction, mRNAs are isolated for library preparation. IVT (unmodified) control library is generated by reverse transcription of mRNA followed by *in vitro* transcription. Libraries are subjected to direct RNA sequencing on the MinION followed by basecalling and alignment, followed by site-specific machine learning (ML) and quantitative Ψ detection. b, Top: example current trace obtained during nanopore sequencing of syn-PSMB2-Ψ synthetic control for the PSMB2 gene that contains a Ψ-site, where each discrete signal fluctuation is associated with presence of a particular RNA k-mer in the pore (k=5). Scheme below illustrates the direction of motor-driven RNA motion through the pore, highlighting the critical positions where the signal is read where the Ψ-centered k-mer is at the pore reader position (pink). Bottom traces show expanded views of the raw signal trace (black) obtained for those sites where Ψ is present in the pore constriction, as well as various Fourier components of the raw signal, used for ML-based Ψ detection. Right: hairline basecalling plot shows the query (top row), with D representing deletions, vs. reference (left column) base calls observed in syn-PSMB2-Ψ reads (n=1,200) at the 5-mer region with Ψ at position 0, where 38.2% U-to-C mismatch error is found (1.8% was found for the syn-PSMB2-U construct). c, Flowchart describing two general approaches for Ψ detection using direct RNA nanopore sequencing. Dashed box represents a U-to-C mismatch error approach to identify Ψ sites, and bottom row represents integration with synthetic mRNAs and machine-learning classification to quantify Ψ occupancy in these sites.

### Selecting a supervised machine learning classifier

We assessed the contribution of upstream features (i.e. features that are related to the presence of Ψ in the protein motor twelve nucleotides upstream) and found that they did not have an impact on model accuracy (**Supplementary Figure S2**). Hence, we continued our analysis with only local Ψ features and removed upstream features, leaving 35 features for ML training. We applied five different supervised ML classifiers for each synthetic construct: logistic regression (LR), gradient boosting (GBC), K-nearest neighbors (KNN), random forest (RF), and support vector machine (SVM). We trained and fit the parameters of each classifier with 75% of the data and assessed its performance with the remaining 25%.

To determine which of the five ML classifiers consistently yields the highest sensitivity and specificity for each construct, we trained each model and evaluated the Receiver Operator Characteristic (ROC) curve and its associated area under the curve (AUC) with the testing dataset (**Fig. 2a-d**, left) The ROC curve was obtained by sweeping the call threshold on the probabilistic output of the models. For the *PSMB2, PRPSAP1*, and *MCM5* synthetic constructs, we observed an AUC equal to or greater than 0.94 for each ML model. Similar results were seen with *MRPS14*, except the AUC for LR and KNN was 0.92 and 0.91, respectively. For the *PSMB2, PRPSAP1, MCM5*, and *MRPS14* synthetic constructs, the RFC and GBC consistently generated equivalent, and highest, AUC results among the five classifiers.

**Figure 2.**
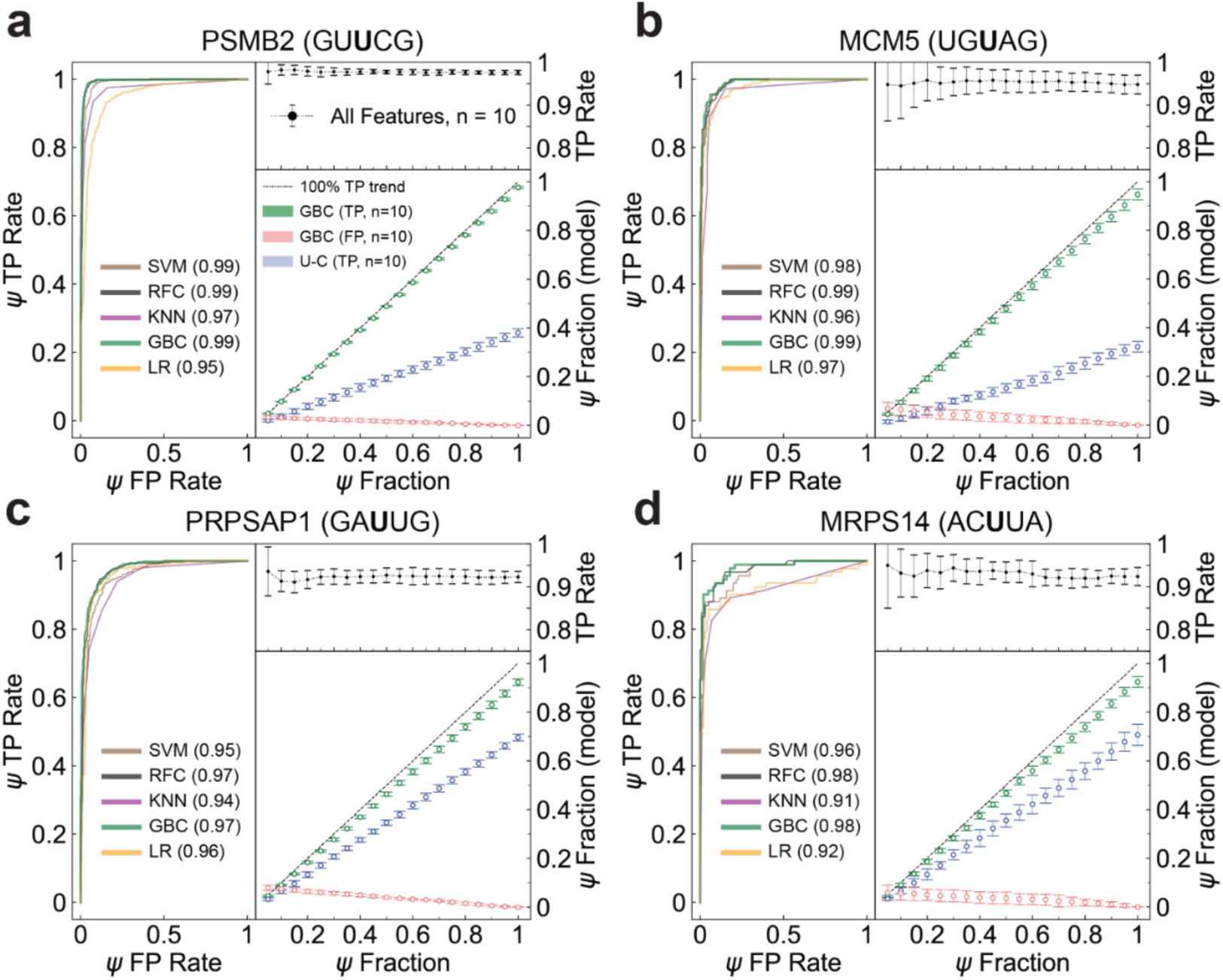
Performance of machine-learning (ML) classification for synthetic Ψ controls. a-d, Machine-learning classification accuracy after training on 35 signal features when Ψ-site in labeled synthetic constructs is present in the pore constriction. Left: Receiver operator characteristic (ROC) curves of five different supervised ML classifiers: support vector machine (SVM), random forest classifier (RFC), K-nearest neighbors (KNN), gradient boosting classifier (GBC), and linear regression (LR), along with their respective area-under-curve values in parentheses. Top right: Mean classification accuracy of Ψ true positive (TP) rate out of total Ψ samples present in the test set (true positives/(true positives+false negatives)) as a function of Ψ fraction in the test set using the GBC classifier (ten models generated for each percentage Ψ with random combinations of data for training and testing). The error bars represent the standard error of the TP rate for those ten models. Bottom right: The mean and standard deviation of TP Ψ prediction by GBC models (the same ten GBC iterations used for the TP rate analysis) divided by total size of the test set is shown in green at each Ψ fraction. Dashed black line represents perfect discrimination of all Ψ and control samples in the test set at each Ψ percent ratio. The mean and standard deviation of false positive (FP) calls by the GBC divided by total size of the test set at each Ψ percent ratio are shown in red. The mean and standard deviation of a U-to-C mismatch classifier divided by the total size of the test at each Ψ percent ratio are shown in blue.

To evaluate which of the two (GBC and RFC) yielded the highest accuracy, we generated 10 random train-test split sets for each classifier and calculated the mean and standard deviation of the model’s accuracy for each set. We found that the GBC classifier consistently had the highest accuracy for all 4 synthetic constructs, with GBC having an accuracy of 0.97±0.00 for *PSMB2*, 0.92±0.01 for *PRPSAP1*, 0.94±0.01 for *MCM5*, 0.94±0.01 for *MRPS14*. In comparison, the next best model, RFC, displayed an accuracy of 0.95±0.00 for *PSMB2*, 0.91±0.01 for *PRPSAP1*, 0.94±0.01 for *MCM5*, and 0.92±0.01 for *MRPS14* (**Supplementary Table S6**). Due to its superior accuracy, we implemented the GBC model for the remainder of our analysis.

### Evaluating the sensitivity and specificity of Ψ detection using the GBC model

To evaluate our capacity to detect Ψ at different occupancies, we generated multiple test sets with different ratios of syn-U and syn-Ψ reads and compared our highest accuracy ML model (GBC) to the U-to-C mismatch error model for the same test sets. In total, we produced 20 different test sets ranging from 5% syn-Ψ (95% syn-U) to 100% syn-Ψ (0% syn-U), in 5% increments. To assess reproducibility, we reshuffled the dataframe 10 times for each synthetic construct, yielding different combinations of training and test sets (see **Methods** for details). In **Fig 2a-d** we show the mean true positive (TP) trend of Ψ classification, calculated by dividing the Ψ calls by the total test set size (green markers). The GBC model performed with a TP accuracy of >90% across all fractions of Ψ for *PRPSAP1, MRPS14, and MCM5*. To determine whether the GBC classifier was overfitting to syn-Ψ reads, we looked at the false positive (FP) trend, which occurs when the model misclassifies syn-U reads as syn-Ψ reads, as a function of Ψ ratio (red markers). As expected, the FP trend had an inverse relationship with syn-Ψ occupancy, <10% for any syn-Ψ ratio for *PSMB2*(0.03), *PRPSAP1*(0.08), *MRPS14*(0.06), and *MCM5*(0.07). As a result, we note that for very low Ψ occupancies (<15%), the model performs poorly in distinguishing Ψ from U.

Finally, we compared the accuracies of the GBC model to the U-to-C mismatch rate Ψ calling by plotting the mean and standard deviation of the U-to-C TP trend (i.e., the ratio of Ψ TP calls to the total size of the test set size) for each artificial syn-Ψ fraction. Notably, the GBC greatly outperforms the U-to-C classifier for all 4 synthetic constructs, despite the fact that U-to-C mismatch rates vary widely from k-mer to k-mer. The U-to-C mismatch rate for PSMB2 was 38% in a 100% syn-Ψ dataset, while the GBC model called 97% of the dataset as Ψ.

What are the most important features that contribute to the accuracy of the GBC model? After training and testing the GBC model, we used scikit-learn’s^26^ *feature importance* tool to obtain the weights for all 35 features, which is an estimate of their relative importance during model fitting (see SI, Tables S7-10). Quality score features were present in the top 10 list for all the synthetic constructs except for *PSMB2*, while Fourier components were only seen in the top 10 list of *MRPS14*. These results demonstrate that the relative importance of individual features is highly dependent on the specific k-mer sequence, as recently suggested by others^19,21^.

### Quantification of site-specific pseudouridine modifications in HeLa cell mRNA

Following the development of an accurate model for Ψ-site detection using our synthetic controls, we applied it to profile Ψ-site occupancies based on three independent direct mRNA nanopore sequencing datasets for HeLa transcriptomes from Tavakoli et al.^22^ The three datasets (D1,D2 and D3) were extracted and filtered using the same filtration steps implemented on the synthetic constructs. Additionally, HeLa IVT reads that aligned to the gene targets were extracted and filtered. After filtration, the 35 features used to fit the GBC model during syn-Ψ training and testing were parsed from each native read and compiled for classification. For each synthetic construct, we generated ten GBC models by fitting each one to a reshuffled synthetic dataset with a split of 85% for training and 15% for testing. Subsequently, each model was invoked onto the HeLa mRNA reads for single-read Ψ prediction, providing a Ψ-quantified output for each gene from D1, D2, and D3 experiments (**Fig. 3a**).

**Figure 3.**
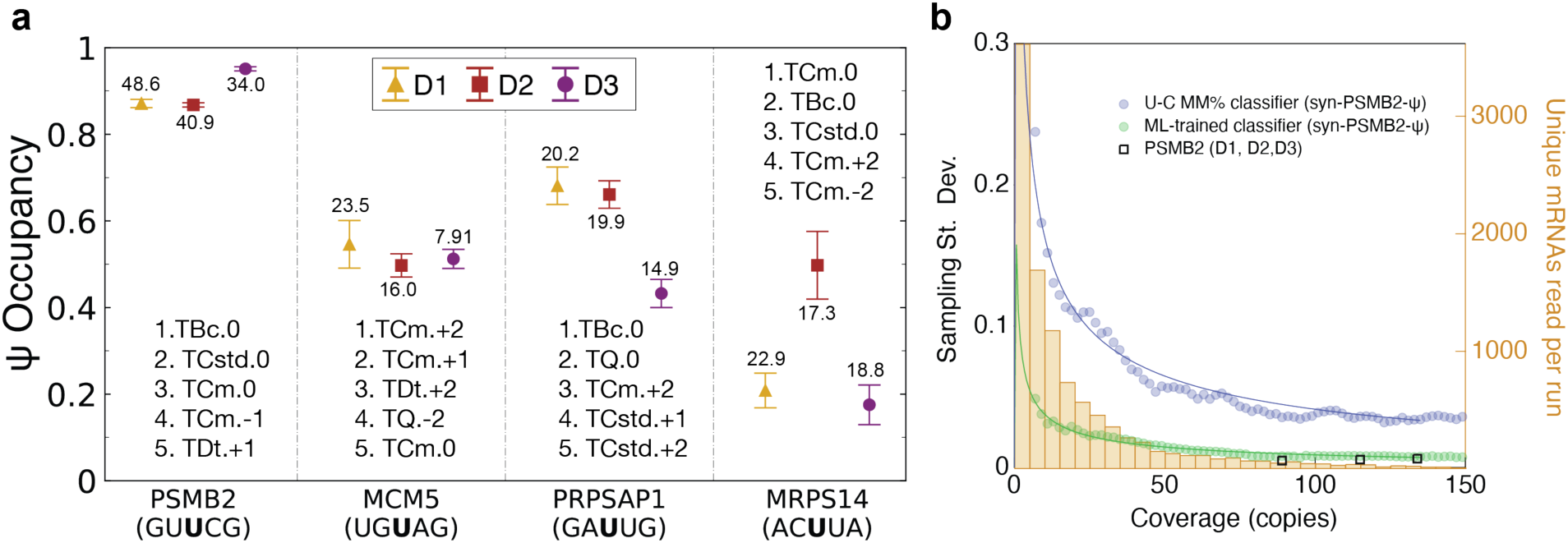
Quantification of Ψ in HeLa cell mRNA with synthetic-trained machine learning models. a, Ψ quantification of Hela mRNA targets (PSMB2, MCM5, PRPSAP1, MRPS14) observed in three independent sequencing libraries, direct 1 (gold) and direct 2 (red), and direct 3 (purple) using a gradient boosting classifier (GBC) trained with the corresponding syn-Ψ and syn-U constructs. The same 35 target features used to train each GBC model were extracted from each Hela mRNA read that passed the necessary filtration stages. The mean and standard deviation illustrate the result of Ψ occupancy called by ten randomly generated GBC models that were all re-corrected using the ratio calibration of true positive and false positive fits observed for each mRNA syn-Ψ construct (green and red trends in Figure 2). The calculated TPM for each gene is annotated next to each marker. The acronyms for the top five weighted features for each replicate-trained GBC model are shown. b, Comparison of Ψ true-positive standard deviation when syn-PSMB2-Ψ reads are predicted with either a U-to-C mismatch classifier (blue) or an GBC (green) as a function of read coverage. Standard deviation of the U-to-C mismatch classifier for each read coverage increment on the x-axis is obtained through resampling reads from the syn-PSMB2-Ψ sample 30 times. A similar approach is used for obtaining GBC Ψ true-positive standard deviation, except that the resampled syn-PSMB2-Ψ reads are only extracted from the test set and not the training set used to build the GBC. The histogram displays the total differential mRNA read coverage captured in the direct 1 library. Square markers indicate actual standard deviations for the three direct RNA sequencing replicates of PSMB2.

Prior to comparing the model-predicted Ψ occupancy for all three direct experiments for each gene, the artificial Ψ TP trend (green markers) and FP trend (red markers), (**Fig. 2a-d** bottom, bottom right) observed during synthetic model training and testing was used to derive a re-correction factor for each gene by summing both together for each Ψ fraction and taking the slope of the linear fit. After reweighing the initial quantified prediction of each model on HeLa mRNA, we observed similar Ψ frequencies (within 10%) across all three independent experiments for *PSMB2 and MCM5* (**Fig. 3a**). For *PSMB2*, we observed a re-corrected mean Ψ-occupancy of 0.89 (D1, D2, and D3). For *MCM5*, the GBC estimated a mean Ψ-occupancy of 0.52. For *PRPSAP1*, the GBC estimated a mean Ψ-occupancy of 0.59. For *MRPS14*, the GBC estimated a mean Ψ-occupancy of 0.29. The read coverage and base mismatch rate per direct experiment for each mRNA are shown in **Supplementary Table S1-4)**. Next to each result in **Fig. 3a** we have annotated the transcripts per million (TPM) count.

To assess the strength of our method in overcoming low-read coverage, we tested and compared the TP rate of our GB classifier with the U-to-C mismatch classifier generated for *PSMB2* as a function of read count with our syn-*PSMB2*-Ψ dataset (**Fig. 3b**). For each read coverage bin, which ranged from n=7 to n=200 in increments of n=2, we resampled from the syn-PSMB2-Ψ dataset multiple times and calculated the TP standard deviation for both the GB and U-to-C mismatch classifiers (green and blue data points, respectively). Compared with the U-to-C mismatch classifier (blue), the standard deviation of the TP rate for our GBC classifier was substantially lower across all read counts. Furthermore, we used the *featureCounts* module from Rsubread^27^ on our direct 1 HeLa mRNA library to corroborate that the majority of mRNA transcripts captured in nanopore sequencing have a relatively low read coverage (**Fig. 3b**, background histogram in gold). Moreover, the standard deviations from ML Ψ quantification results for all *PSMB2* direct RNA sequencing data (**Fig. 3a**) are in agreement with the fit of the TP rate standard deviation versus read coverage for the *PSMB2*-trained GBC (**Fig. 3b**).

## Discussion

It has been previously established that U-to-C mismatch error may be used to identify sites of pseudouridine modification^18–22^. However, based on the synthetic controls established in Tavakoli et al., the variable U-to-C mismatch rate for the Ψ-modified synthetic controls demonstrates that this method is not quantitative, and highly dependent on the sequence context. The Guppy (3.2.10) basecaller was trained on a heterogenous population of RNAs containing a majority of canonical nucleotides, but also containing modified nucleotides. Since this basecaller was not trained on k-mers that exclusively contain Ψ-sites, mismatch errors are inconsistent (**Fig. 1b**), requiring re-training in the right sequence context in order to accurately distinguish Ψ from canonical U.

To determine what combination of features can enhance Ψ discrimination, we extracted from the sequencing data a total of 60 raw signal features and found that 35 local features were critical for Ψ discrimination (**Supplementary Table S5**). Upstream features corresponding to presence of the suspect Ψ-site in the protein motor (12 nucleotides upstream from the Ψ-site in the pore) were considered because of a recent report^21^ that showed Ψ modifications with an adjacent 5’ guanosine (G) can induce distinct pauses in motor protein steps. However, Stephenson et al.^28^ showed that G-rich RNA sequences can also stall the motor protein, making it a less reliable parameter under circumstances where Ψ is near or on a polyG region. We therefore excluded these features because these did not provide a noticeable boost in accuracy (**Supplementary Figure S2**).

Five different supervised ML models were tested for each synthetic construct, and we found that GBC consistently provided the highest classification accuracy for every synthetic replicate. Conversely, previous algorithms have used KNN for Ψ quantification which we observed to have the lowest AUC and classification accuracy (**Supplementary Table S6**). This may be attributed to the high dimensional feature space (35 dimensions) of our training data, which is not suitable for KNN, and that was previously trained on a 6-dimensional data set based on the quality scores of three bases and the current mean of three 5-mers (−1, 0 [U/Ψ], +1)^19^. GBC accuracies were high across different constructs, with mean TP rates (TP/(TP+FP)) >0.9 for *PSMB2* (GU**U**CG), *PRPSAP1* (GA**U**UG), *MCM5* (UG**U**AG), and *MRPS14* (AC**U**UA) across all concentration increments from 5% - 100% (**Fig 2**, top right). Evaluating the top five weighted features for the trained models (see **Supplementary Information**, page 16) generated for each construct using scikit-learn’s *feature importance* revealed that not a single feature was retained across all synthetic constructs. We observed a variable degree of separation and difference in correlation among the top five weighted features between syn-Ψ and syn-U with respect to each construct (**Supplementary S4-S11**). Moreover, we found signal features that correspond to 5-mers with Ψ in the +2 and –2 to be among the top five fitting parameters for MCM5 (UG**U**AG), MRPS14 (AC**U**UA), and PRPSAP1 (GA**U**UG), with current mean of the +2 5-mer having the highest weight for MCM5.These results further highlight the critical need to train models that consider the sequence context neighboring the modified site, which should also ensure all 5-mers bearing Ψ in every position (−2, -1, 0, +1, +2) to be sequenced without the influence of any neighboring Ψ modifications.

Finally, we applied our computational engine to HeLa mRNA reads that aligned to the corresponding gene from three biological replicates. Remarkably, the Ψ occupancy called by the GBC model was similar for all three direct experiments for *PSMB2 (chr1: 35603333)* and *MCM5 (chr22: 35424407)*. Based on the functions of these genes, *PSMB2* is a component of the 20S core proteasome complex that degrades most intracellular proteins, while *MCM5* is involved in the initiation of DNA replication during mitosis. For the *MRPS14 (chr1: 175014468)*, the GBC predicted similar Ψ-occupancy across D1 and D3, while there was a noticeable increase in the Ψ-occupancy at D2 (**Fig 3a**). The GBC trained for *PRPSAP1(chr17: 76311411)* estimated a similar Ψ-occupancy for D1 and D2 at ∼65%, while D3 had a lower Ψ-occupancy at ∼45%. The computational engine we developed here achieves the first quantitative Ψ occupancy measurements in human mRNAs from direct RNA sequencing data. This 2-step engine first integrates endogenous transcriptome data to identify putative sites *de novo* via specific U-to-C basecalling errors^22^), and then quantifies the Ψ occupancy at a given site using ML models that are trained on sequence-specific synthetic mRNA standards. Our application of supervised ML models reveals that for each Ψ-site studied, different signal parameters are required to maximize Ψ classification accuracy. Applying our models resulted in quantification of these Ψ-sites with a much higher accuracy than U-to-C basecalling errors provide. We show that this improved quantification ability of our engine is particularly critical for low-abundance mRNAs, for which typical mRNA coverage are low for a MinION run (<10). Additional synthetic controls for validated Ψ-sites applied in combination with our new computational engine, would enable us to profile patterns of Ψ mRNA modification with high accuracy from minION direct RNA sequencing libraries.

## Methods

### Alignment

After basecalling with guppy (3.2.10), fastq reads that passed the default ONT filtration stage (>Q7) were aligned to the synthetic reference using minimap 2 (2.17) with the option ‘‘-ax map-ont -un -k15’’. The sam file was converted to bam using samtools (1.10). Bam files were sorted by “samtools sort” and indexed using “samtools index” and visualized using IGV (2.8.13). Finally, a bam file was prepared for each synthetic construct by slicing out the corresponding reads from the original bam file using “samtools view -h -Sb”.

### Filtration

After alignment, a second-stage filtration step was implemented to remove reads that were truncated near the site of modification. For the synthetic replicates, Ψ was located on position 511 for all four transcripts. Each read was scanned for the position of Ψ, the 7 bases upstream (3’) from Ψ, and the 7 basecalls downstream (5’) from Ψ, denoting this region as the 15-mer target segment. Next, using Rsamtools (3.6.0), we set the filter pass conditions to only retain reads with a mapping quality score of 50. Additionally, each read was required to have no more than three deletions within its 15-mer target segment. Finally, reads with one or more insertions in the 15mer target segment were filtered out. Reads that were retained after this stage were passed onto the next stage for feature extraction.

### Feature extraction

Basecalls and quality scores were extracted with Rsamtools (3.6.0). Current data used to prepare signal features was extracted using nanopolish *eventalign*.

### Data preprocessing

For each construct, features from syn-Ψ and syn-U reads were labeled and combined into one dataframe. We used the scikit-learn python library (1.0.2) for data preprocessing and model training and testing. For each replicate, the dataset was resampled to contain an equal sample size of both unmodified and Ψ modified transcripts. The Ψ modified data was the limiting factor for all four targets. Next, the dataset underwent a 75/25 split, where 75% of the reads were randomly binned into the training set and the remaining 25% went into the test set. The features in the training set were normalized using the scikit-learn’s *StandardScaler* function, where the mean was centered around 0 and the first standard deviation was +/-1. The normalization parameters were then used to scale the features in the test set.

### Model training/testing/evaluation

The five supervised ML models (support vector machine, logistic regression, random forest, and k-nearest neighbors) were imported from scikit-learn. Every model was trained and tested with each construct-specific dataframe. The accuracy and reproducibility of the models were assessed through multiple training and testing iterations (n=10), where for every model generation, the dataframe was reshuffled in order to produce a new 75-25 split. ROC plots were made with scikit-learn’s *plot_roc_curve*. Model accuracy as a function of Ψ concentration (**Fig 2**) was acquired from 10 different models that were generated with a balanced training set and subsequently implemented on test sets that varied in Ψ:U ratio. The original test set was resampled to get the desired Ψ ratio.

### HeLa mRNA classification and analysis

The same features from synthetic reads used for model generation were extracted and prepared from native reads. Prior to Ψ quantification of native reads, the GBC model was trained on synthetic data corresponding to the native reads with an 85-15 split. Native data was normalized with the same parameters used to scale the testing data. Next, the model classified every native read as Ψ or unmodified. This process was repeated ten times, with each model having a different train-test split. Finally, the reported Ψ percentage present in the native reads was recorrected with the addition of the model’s average false positive and true positive values observed from the analysis that tested model accuracy as a function of Ψ occupancy (**Fig 2**).

## Code availability

Scripts for all analyses presented in this paper, including all data extraction, processing, and graphing steps are freely accessible at https://github.com/wanunulab/psiquant.

## Data availability

All raw and processed data used to generate figures and representative images presented in this paper are available at https://www.biorxiv.org/content/10.1101/2021.11.03.467190v1.

## Statistical analysis

All experiments were performed in multiple, independent experiments, as indicated in the figure legends. All statistics and tests are described fully in the text or figure legend.

## ACKNOWLEDGMENTS

We acknowledge Dr. Miten Jain for helpful advice with data preparation for processing. The authors acknowledge generous support through an Opportunity Fund by the Technology Development Coordinating Center at Jackson Laboratories (NHGRI federal award no. U24HG011735).

