## Supplementary Figures for "Messenger-RNA Modification Standards and Machine Learning Models Facilitate Absolute Site-Specific Pseudouridine Quantification"

**Table S1:** Sample size, U-to-C mismatch rate, and U-to-N mismatch rate of **MCM5** synthetic and native mRNA used in this study

| Experiment | N Reads | U-to-C MM% | U-to-N MM% |
|---|---|---|---|
| Syn-U | 7890 | 5.62% | 7.17% |
| syn-ψ | 687 | 33.45% | 52.31% |
| Direct 1 | 51 | 56.86% | 58.82% |
| Direct 2 | 69 | 47.82% | 50.72% |
| Direct 3 | 57 | 50.87% | 52.63% |
| IVT | 87 | 5.74% | 6.90% |

**Table S2:** Sample size, U-to-C mismatch rate, and U-to-N mismatch rate of **MRPS14** synthetic and native mRNA used in this study

| Experiment | N Reads | U-to-C MM% | U-to-N MM% |
|---|---|---|---|
| Syn-U | 2990 | 2.94% | 9.53% |
| syn-ψ | 385 | 69.35% | 79.22% |
| Direct 1 | 23 | 26.08% | 30.43% |
| Direct 2 | 21 | 57.14% | 61.90% |
| Direct 3 | 27 | 11.11% | 14.81% |
| IVT | 123 | 5.70% | 5.70% |

**Table S3:** Sample size, U-to-C mismatch rate, and U-to-N mismatch rate of **PRPSAP1** synthetic and native mRNA used in this study

| Experiment | N Reads | U-to-C MM% | U-to-N MM% |
|---|---|---|---|
| Syn-U | 3671 | 3.02% | 4.00% |
| Syn-ψ | 2184 | 69.64% | 74.54% |
| Direct 1 | 22 | 40.90% | 40.90% |
| Direct 2 | 24 | 50.00% | 50.00% |
| Direct 3 | 11 | 18.18% | 18.18% |
| IVT | 54 | 0.00% | 1.85% |

**Table S4:** Sample size, U-to-C mismatch rate, and U-to-N mismatch rate of **PSMB2** synthetic and native mRNA used in this study

| Experiment | N Reads | U-to-C MM% | U-to-N MM% |
|---|---|---|---|
| Syn-U | 6201 | 1.79% | 2.67% |
| Syn-ψ | 3020 | 38.17% | 47.54% |
| Direct 1 | 113 | 72.56% | 75.22% |
| Direct 2 | 135 | 79.56% | 82.22% |
| Direct 3 | 89 | 86.51% | 88.76% |
| IVT | 303 | 1.65% | 4.62% |

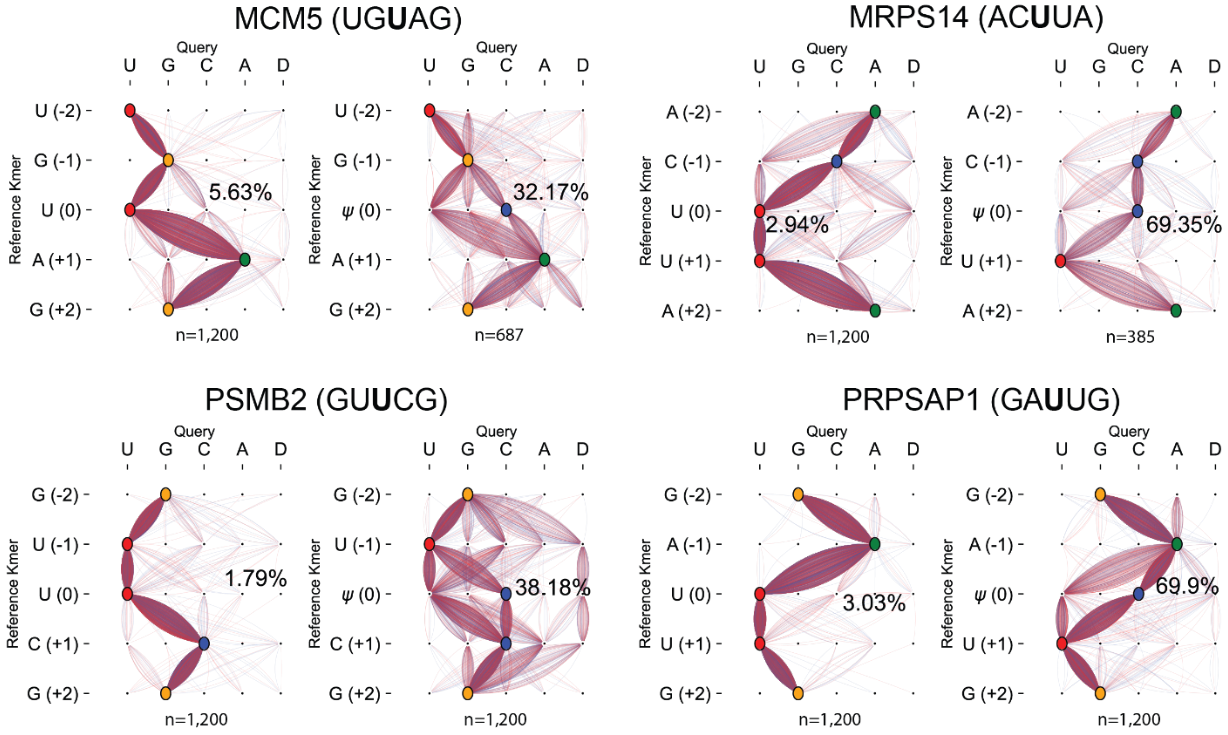

**Figure S1**: Basecaller (Guppy) performance on Kmer (k=5) sequence region where the ψ-modified or canonical U nucleotide is present in position 0. The correct K-mer sequence is shownon the vertical axis (Reference Kmer), while the basecalls made (Query) as the K-mer is traversed is shown. Basecalls of unmodified syn-U reads are shown on the left while the modified syn- ψ reads are shown on the right. The U-to-C mismatch rate (%) for each set of synthetic reads is annotated.

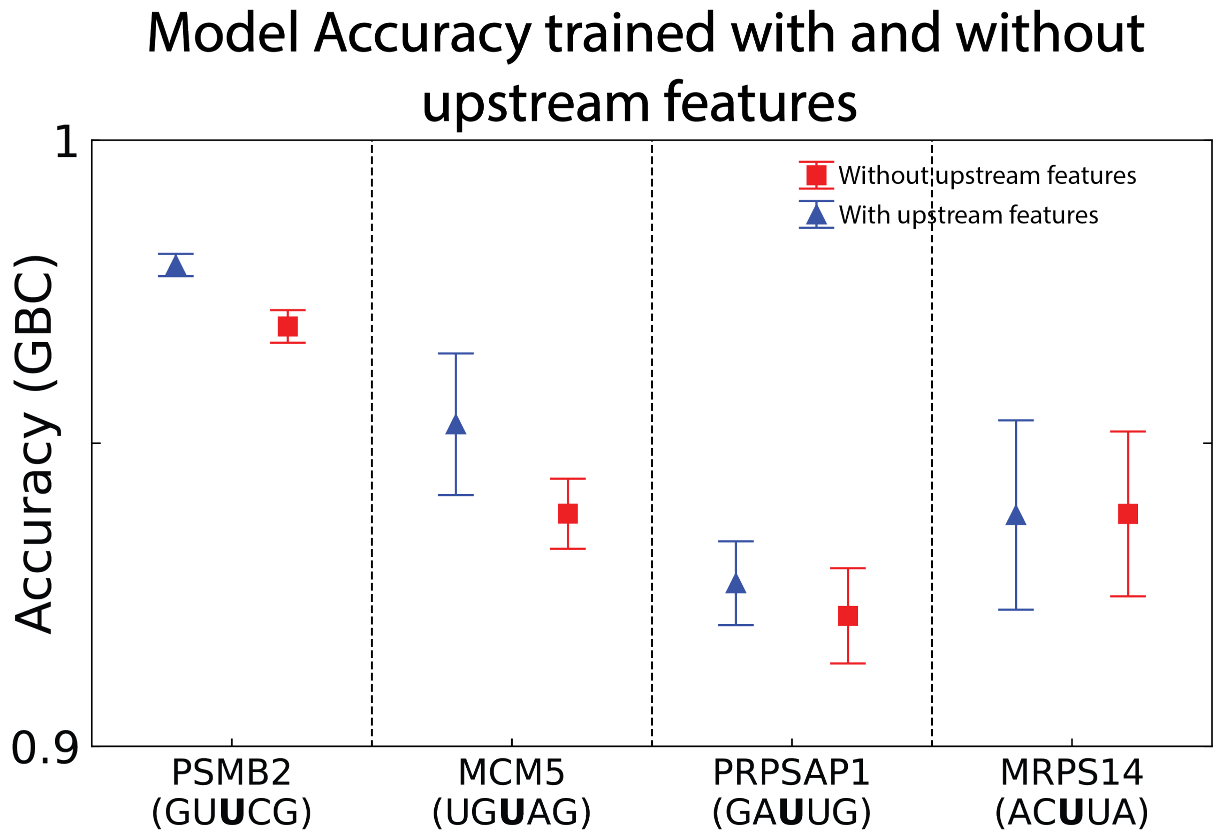

**Figure S2**: Comparison of GBC model accuracy trained with features parsed from synthetic transcripts that correspond to the presence of the modified pseudouridine/unmodified uridine nucleotide present inside the constriction of the pore (35 total features, red) versus the same set of features with the addition of upstream features that correspond to the presence of the modified/unmodified pseudouridine nucleotide present inside the helicase motor protein situated on top of the pore.

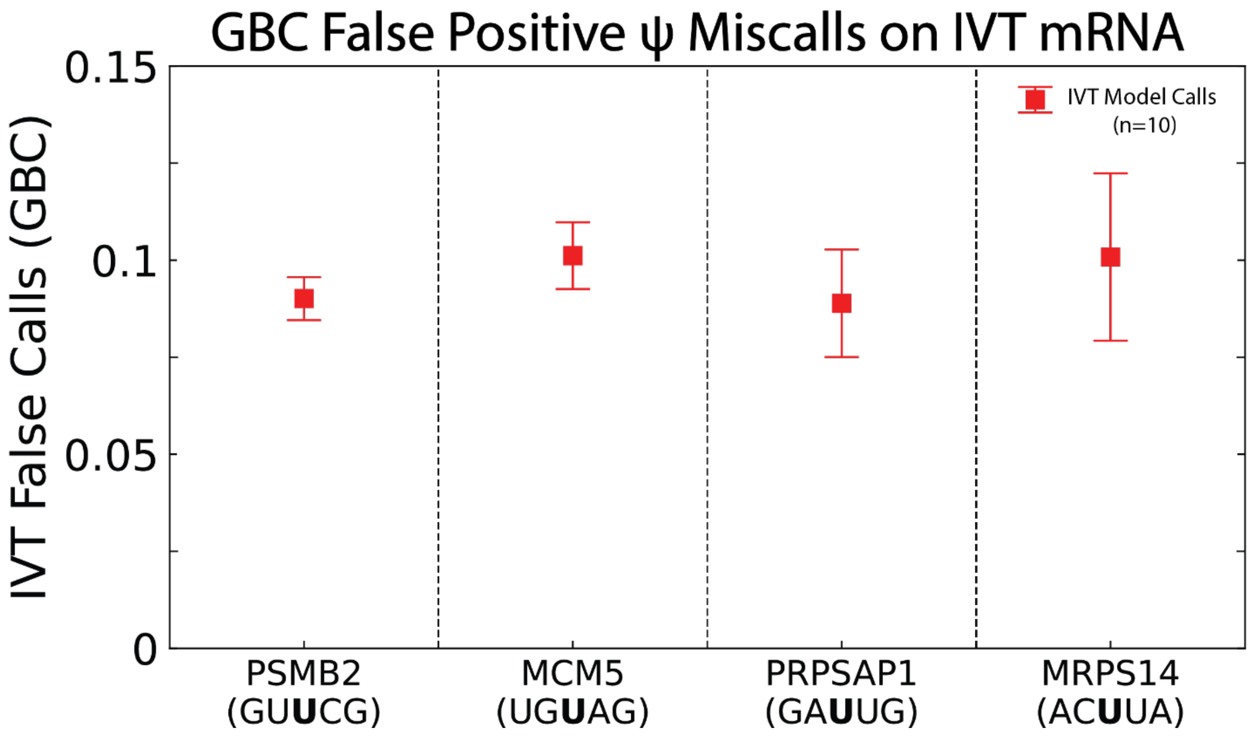

**Figure S3**: Pseudouridine false-positive miscalls on direct *in vitro* transcribed (IVT) mRNA (unmodified) by each GB classifier trained with synthetic transcripts for all four genes. The respective GBC model generated for each gene was retrained on a reshuffled synthetic dataset and subsequently applied for IVT classification ten times.

**Table S5:** Description key containing each feature (including upstream features) extracted and used for fitting machine-learning models. For base call and quality score features, the positions go from 5' to 3' (-2 to +2), while the signal feature positions (Kmer frames) go from 3' to 5' (-2 to +2). Note that the Kmer position is based on which nucleotide is located in the center of the of the 5-mer relative to the position of the modified/unmodified nucleotide.

| Program Feature name | Acronym | Full Description |
|---|---|---|
| Bn2 | TBc.-2 | Base Call at position -2 |
| Bn1 | TBc.-1 | Base Call at position -1 |
| B0 | TBc.0 | Base Call at position 0 (ψ) |
| Bp1 | TBc.+1 | Base Call at position +1 |
| Bp2 | TBc.+2 | Base Call at position +2 |
| Qn2 | TQ.-2 | Quality Score of base at position -2 |
| Qn1 | TQ.-1 | Quality Score of base at position -1 |
| Q0 | TQ.0 | Quality Score of base at position 0 |
| Qp1 | TQ.+1 | Quality Score of base at position +1 |
| Qp2 | TQ.+2 | Quality Score of base at position +2 |
| current_mean_n2 | TCm.-2 | Current mean of signal when base in -2 position is present in the center of the pore constriction |
| current_mean_n1 | TCm.-1 | Current mean of signal when base in -1 position is present in the center of the pore constriction |

| | | |
|---|---|---|
| current_mean_0 | TCm.0 | Current mean of signal when base in 0 position is present in the center of the pore constriction |
| current_mean_p1 | TCm.+1 | Current mean of signal when base in +1 position is present in the center of the pore constriction |
| current_mean_p2 | TCm.+2 | Current mean of signal when base in +2 position is present in the center of the pore constriction |
| current_std_n2 | TCstd.-2 | Current standard deviation of signal when base in -2 position is present in the center of the pore constriction |
| current_std_n1 | TCstd.-1 | Current standard deviation of signal when base in -1 position is present in the center of the pore constriction |
| current_std_0 | TCstd.0 | Current standard deviation of signal when base in 0 position is present in the center of the pore constriction |
| current_std_p1 | TCstd.+1 | Current standard deviation of signal when base in +1 position is present in the center of the pore constriction |
| current_std_p2 | TCstd.+2 | Current standard deviation of signal when base in +2 position is present in the center of the pore constriction |
| upstream_current_mean_n2 | UCm.-2 | Current mean of signal when the base 14 nucleotides upstream from ψ/U is present in the center of the pore constriction |
| upstream_current_mean_n1 | UCm.-1 | Current mean of signal when the base 13 nucleotides upstream from ψ/U is present in the center of the pore constriction |
| upstream_current_mean_0 | UCm.0 | Current mean of signal when the base 12 nucleotides upstream from ψ/U is present in the center of the pore constriction |
| upstream_current_mean_p1 | UCm.+1 | Current mean of signal when the base 11 nucleotides upstream from ψ/U is present in the center of the pore constriction |
| upstream_current_mean_p2 | UCm.+2 | Current mean of signal when the base 10 nucleotides upstream from ψ/U is present in the center of the pore constriction |
| upstream_current_std_n2 | UCstd.-2 | Current standard deviation of signal when the base 14 nucleotides upstream from ψ/U is present in the center of the pore constriction |
| upstream_current_std_n1 | UCstd.-1 | Current standard deviation of signal when the base 13 nucleotides upstream from ψ/U is present in the center of the pore constriction |
| upstream_current_std_0 | UCstd.0 | Current standard deviation of signal when the base 12 nucleotides upstream from ψ/U is present in the center of the pore constriction |
| upstream_current_std_p1 | UCstd.+1 | Current standard deviation of signal when the base 11 nucleotides upstream from ψ/U is present in the center of the pore constriction |
| upstream_current_std_p2 | UCstd.+2 | Current standard deviation of signal when the base 10 nucleotides upstream from ψ/U is present in the center of the pore constriction |
| dwell_time_n2 | TDt.-2 | Dwell time of signal when base in -2 position is present in the center of the pore constriction |
| dwell_time_n1 | TDt.-1 | Dwell time of signal when base in -1 position is present in the center of the pore constriction |

| | | |
|---|---|---|
| dwell_time_0 | TDt.0 | Dwell time of signal when base in 0 position is present in the center of the pore constriction |
| dwell_time_p1 | TDt.+1 | Dwell time of signal when base in +1 position is present in the center of the pore constriction |
| dwell_time_p2 | TDt.+2 | Dwell time of signal when base in +2 position is present in the center of the pore constriction |
| upstream_dwell_time_n2 | UDt.-2 | Dwell time of signal when the base 14 nucleotides upstream from ψ/U is present in the center of the pore constriction |
| upstream_dwell_time_n1 | UDt.-1 | Dwell time of signal when the base 13 nucleotides upstream from ψ/U is present in the center of the pore constriction |
| upstream_dwell_time_0 | UDt.0 | Dwell time of signal when the base 12 nucleotides upstream from ψ/U is present in the center of the pore constriction |
| upstream_dwell_time_p1 | UDt.+1 | Dwell time of signal when the base 11 nucleotides upstream from ψ/U is present in the center of the pore constriction |
| upstream_dwell_time_p2 | UDt.+2 | Dwell time of signal when the base 10 nucleotides upstream from ψ/U is present in the center of the pore constriction |
| ff1_target_n2 | TFC2.-2 | 2nd Fourier Coefficient of signal when base in -2 position is present in the center of the pore constriction |
| ff2_target_n2 | TFC3.-2 | 3rd Fourier Coefficient of signal when base in -2 position is present in the center of the pore constriction |
| ff1_target_n1 | TFC2.-1 | 2nd Fourier Coefficient of signal when base in -1 position is present in the center of the pore constriction |
| ff2_target_n1 | TFC3.-1 | 3rd Fourier Coefficient of signal when base in -1 position is present in the center of the pore constriction |
| ff1_target_0 | TFC2.0 | 2nd Fourier Coefficient of signal when base in 0 position is present in the center of the pore constriction |
| ff2_target_0 | TFC3.0 | 3rd Fourier Coefficient of signal when base in 0 position is present in the center of the pore constriction |
| ff1_target_p1 | TFC2.+1 | 2nd Fourier Coefficient of signal when base in +1 position is present in the center of the pore constriction |
| ff2_target_p1 | TFC3.+1 | 3rd Fourier Coefficient of signal when base in +1 position is present in the center of the pore constriction |
| ff1_target_p2 | TFC2.+2 | 2nd Fourier Coefficient of signal when base in +2 position is present in the center of the pore constriction |
| ff2_target_p2 | TFC3.+2 | 3rd Fourier Coefficient of signal when base in +2 position is present in the center of the pore constriction |
| ff1_upstream_n2 | UFC2.-2 | 2nd Fourier Coefficient of signal when the base 14 nucleotides upstream from ψ/U is present in the center of the pore constriction |
| ff2_upstream_n2 | UFC3.-2 | 3rd Fourier Coefficient of signal when the base 14 nucleotides upstream from ψ/U is present in the center of the pore constriction |
| ff1_upstream_n1 | UFC2.-1 | 2nd Fourier Coefficient of signal when the base 13 nucleotides upstream from ψ/U is present in the center of the pore constriction |

| | | |
|---|---|---|
| ff2_upstream_n1 | UFC3.-1 | 3rd Fourier Coefficient of signal when the base 13 nucleotides upstream from ψ/U is present in the center of the pore constriction |
| ff1_upstream_0 | UFC2.0 | 2nd Fourier Coefficient of signal when the base 12 nucleotides upstream from /U is present in the center of the pore constriction |
| ff2_upstream_0 | UFC3.0 | 3rd Fourier Coefficient of signal when the base 12 nucleotides upstream from ψ/U is present in the center of the pore constriction |
| ff1_upstream_p1 | UFC2.+1 | 2nd Fourier Coefficient of signal when the base 11 nucleotides upstream from ψ/U is present in the center of the pore constriction |
| ff2_upstream_p1 | UFC3.+1 | 3rd Fourier Coefficient of signal when the base 11 nucleotides upstream from ψ/U is present in the center of the pore constriction |
| ff1_upstream_p2 | UFC2.+2 | 2nd Fourier Coefficient of signal when the base 10 nucleotides upstream from ψ/U is present in the center of the pore constriction |
| ff2_upstream_p2 | UFC3.+2 | 3rd Fourier Coefficient of signal when the base 10 nucleotides upstream from ψ/U is present in the center of the pore constriction |

**Table S6:** Classification accuracy (mean±standard deviation) for each machine learning algorithm with respect to each replicate gene, where each algorithm was trained and tested ten times with synthetic reads used in this study.

| Gene | Gradient Boosting | Random Forest | Logistic Regression | Support Vector Machine | K-Nearest Neighbors |
|---|---|---|---|---|---|
| MCM5 | 0.94±0.006 | 0.94±0.011 | 0.91±0.012 | 0.93±0.011 | 0.87±0.012 |
| MRPS14 | 0.94±0.024 | 0.92±0.014 | 0.90±0.017 | 0.92±0.015 | 0.88±0.021 |
| PRPSAP1 | 0.92±0.008 | 0.91±0.005 | 0.91±0.009 | 0.89±0.008 | 0.87±0.010 |
| PSMB2 | 0.97±0.003 | 0.95±0.003 | 0.89±0.007 | 0.95±0.003 | 0.92±0.005 |

**Table S7:** Normalized weight of ten most important features in GBC model trained with syn-MCM5-ψ and syn-MCM5-U transcripts.

| Feature | Weight |
|---|---|
| current_mean_p2 | 0.348 |
| current_mean_p1 | 0.133 |
| dwell_time_p2 | 0.112 |
| Qn2 | 0.078 |
| current_mean_0 | 0.053 |
| current_mean_n2 | 0.048 |
| B0 | 0.035 |
| current_mean_n1 | 0.030 |
| current_std_p2 | 0.026 |
| current_std_p1 | 0.020 |

**Table S8:** Normalized weight of ten most important features in GBC model trained with syn-MRPS14-ψ and syn-MRPS14-U transcripts.

| Feature | Weight |
|---|---|
| current_mean_0 | 0.548 |
| B0 | 0.148 |
| current_std_0 | 0.034 |
| current_mean_p2 | 0.033 |
| current_std_n2 | 0.029 |
| ff1_target_n2 | 0.020 |
| current_std_n1 | 0.020 |
| Qn1 | 0.018 |
| dwell_time_0 | 0.014 |
| Q0 | 0.011 |

**Table S9:** Normalized weight of ten most important features in GBC model trained with syn-PRPSAP1-ψ and syn-PRPSAP1-U transcripts.

| Feature | Weight |
|---|---|
| B0 | 0.548 |
| Q0 | 0.148 |
| current_mean_p2 | 0.034 |
| current_std_p1 | 0.033 |
| current_std_p2 | 0.029 |
| current_mean_0 | 0.020 |
| Bp1 | 0.020 |
| current_mean_n1 | 0.018 |
| dwell_time_0 | 0.014 |
| Qp2 | 0.011 |

**Table S10:** Normalized weight of ten most important features in GBC model trained with syn-PSMB2-ψ and syn-PSMB2-U transcripts.

| Feature | Weight |
|---|---|
| B0 | 0.249 |
| current_std_0 | 0.220 |
| current_mean_0 | 0.217 |
| current_mean_n1 | 0.089 |
| dwell_time_p1 | 0.039 |
| current_mean_p1 | 0.035 |
| current_std_p2 | 0.023 |
| current_std_p1 | 0.022 |
| current_std_n1 | 0.014 |
| current_mean_n2 | 0.013 |

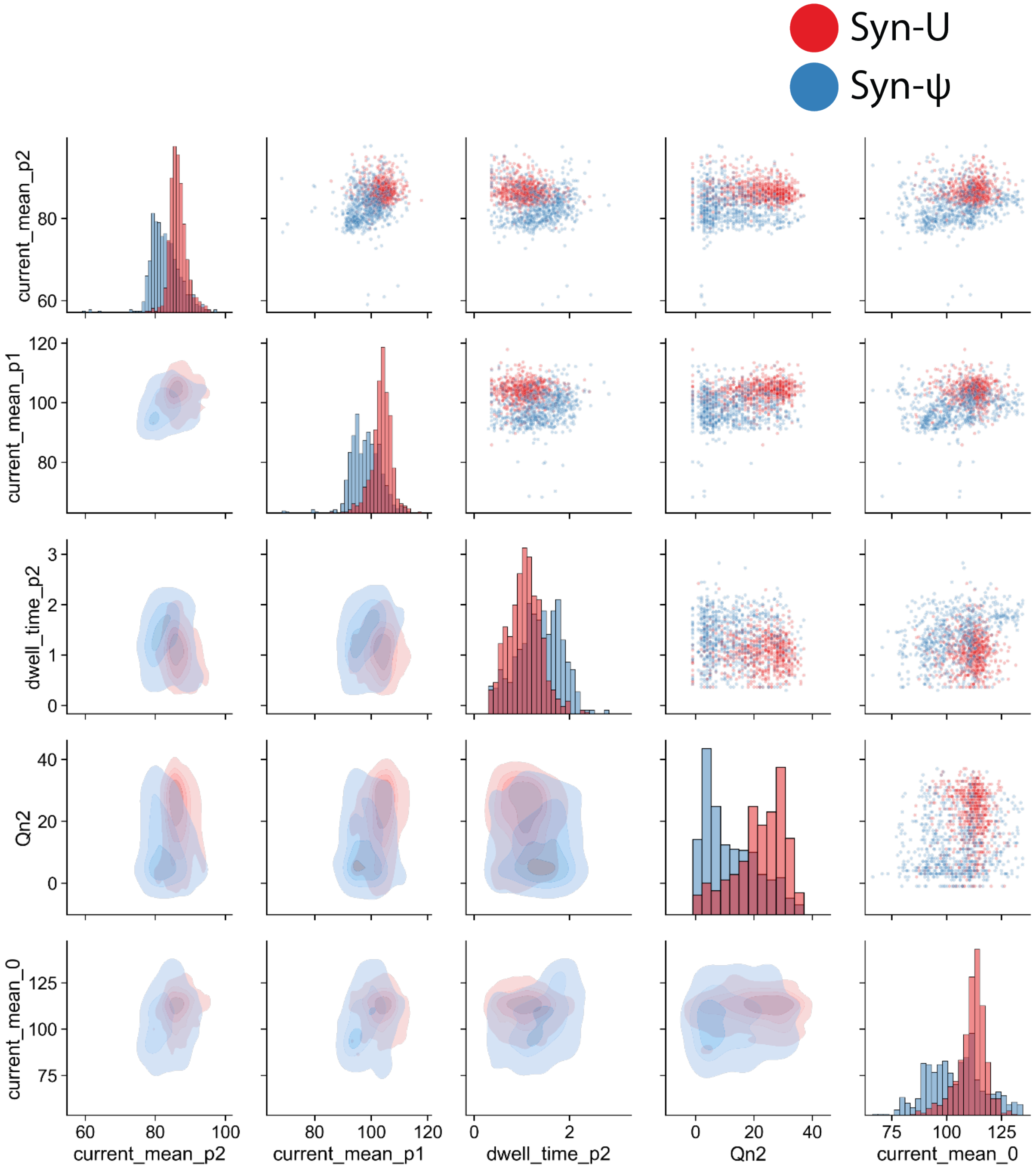

**Figure S4**: Separation of top 5 weighted GBC model features in syn-MCM5-ψ (UG**ψ**AG, blue) syn-MCM5-U (UG**U**AG, red) synthetic training data. Features are compared with 2D scatter plots (upper-half), histogram distribution for each feature (diagonal), and contour plots (lower-half).

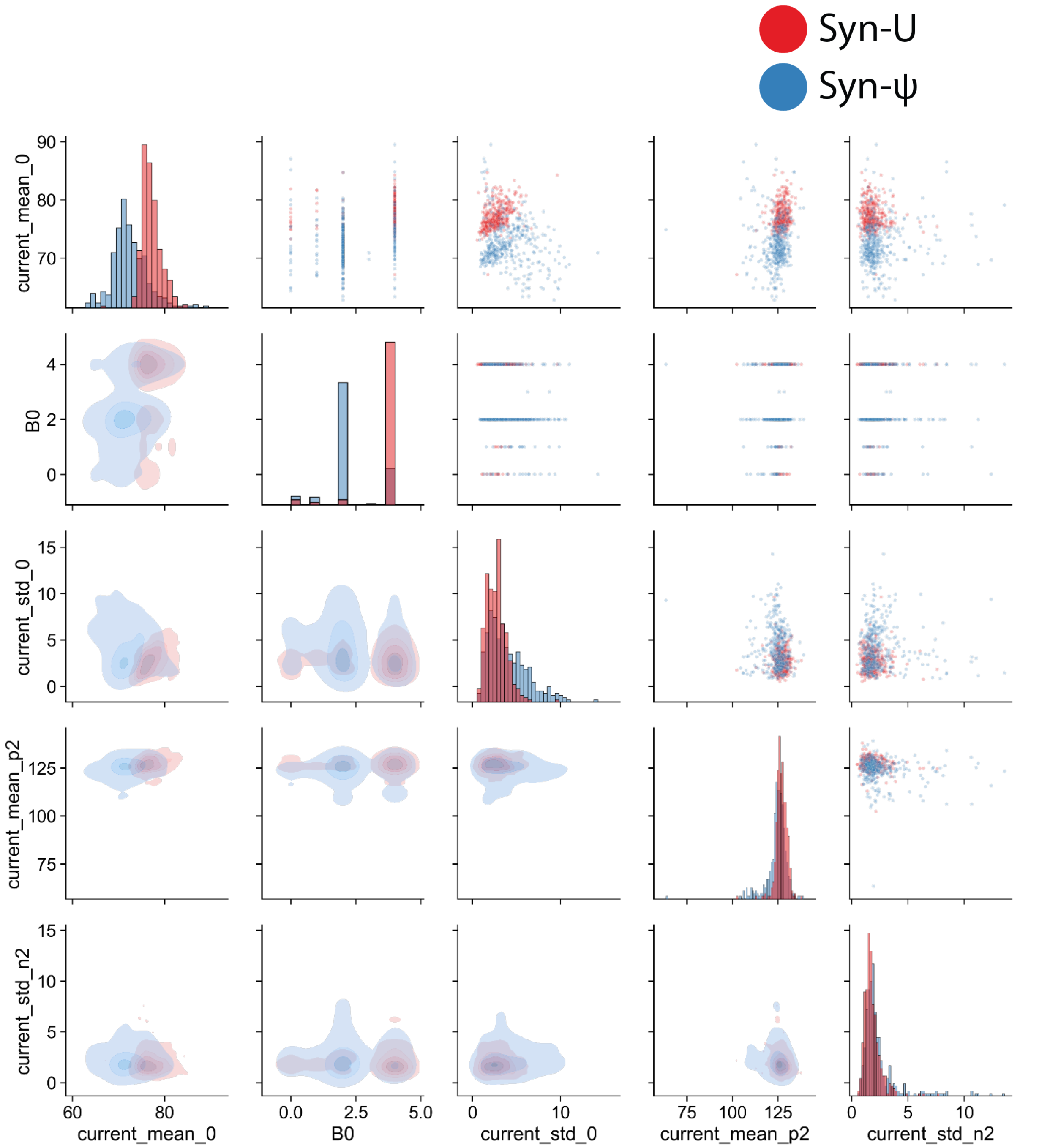

**Figure S5**: Separation of top 5 weighted GBC model features in syn-MRPS14-ψ (AC**ψ**UA, blue) syn-MRPS14-U (AC**U**UA, red) synthetic training data. Features are compared with 2D scatter plots (upper-half), histogram distribution for each feature (diagonal), and contour plots (lower-half).

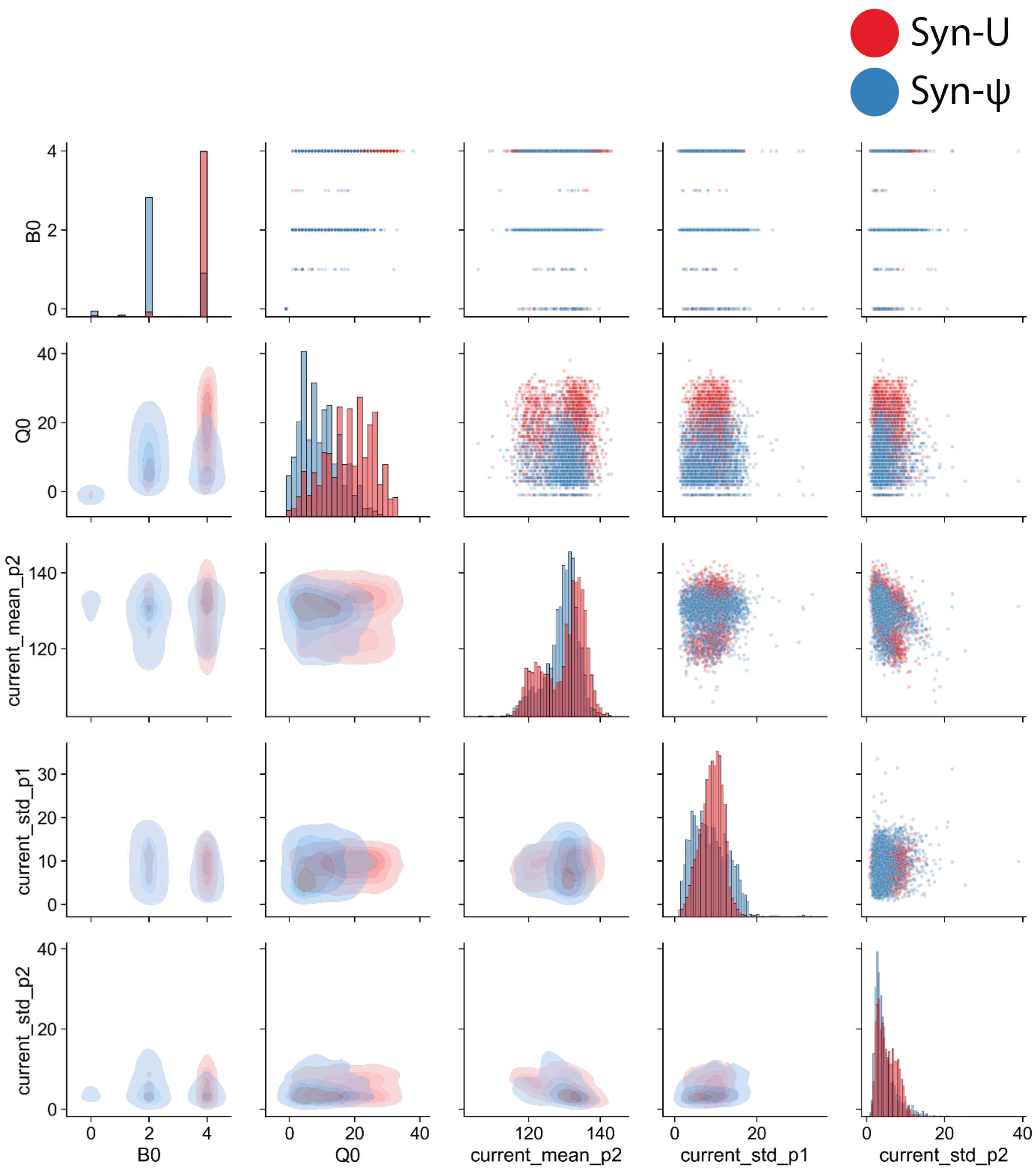

**Figure S6:** Separation of top 5 weighted GBC model features in syn-PRPSAP1-ψ (GA**ψ**UG, blue) syn-PRPSAP1-U (GA**U**UG, red) synthetic training data. Features are compared with 2D scatter plots (upper-half), histogram distribution for each feature (diagonal), and contour plots (lower-half).

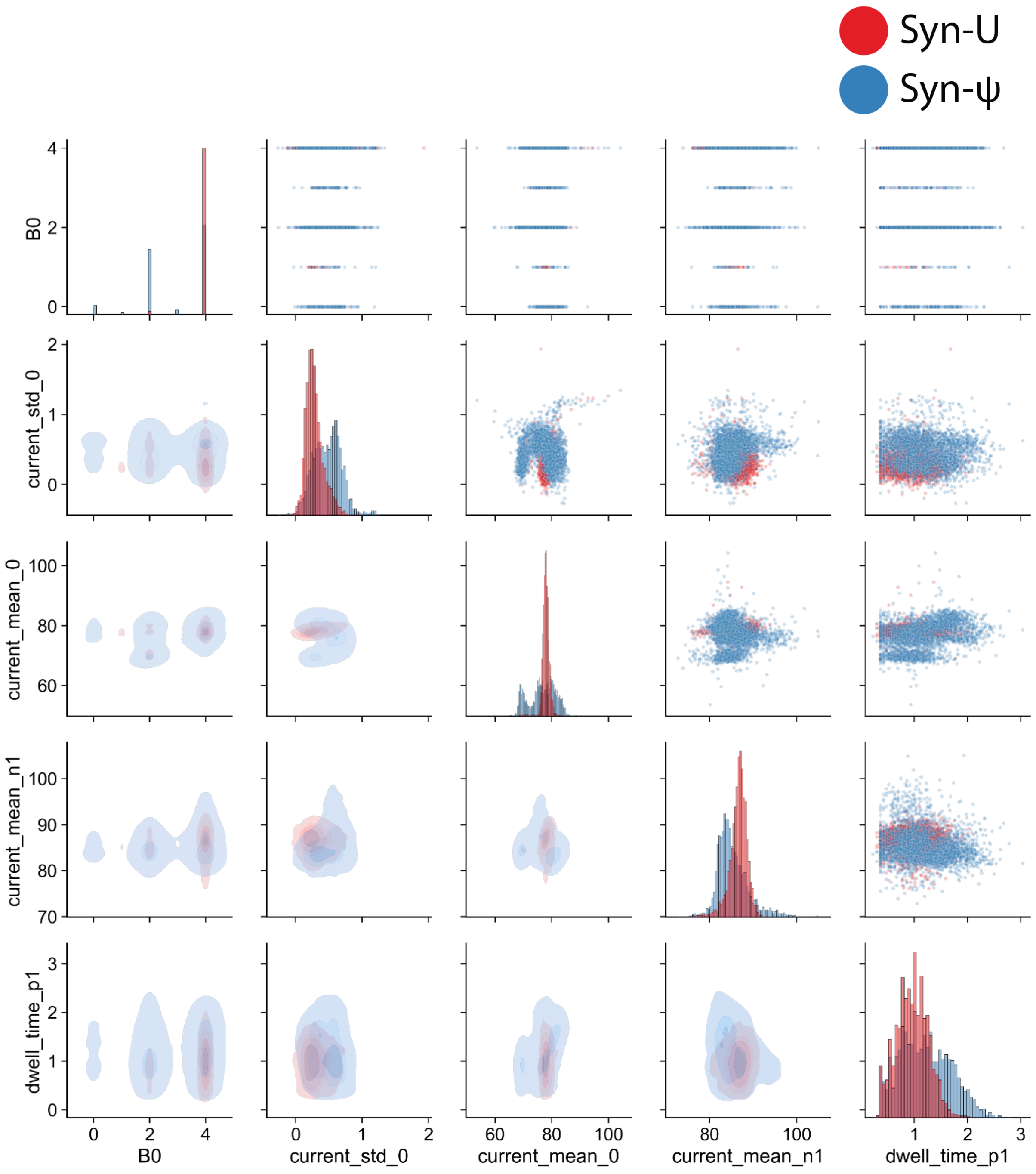

**Figure S7**: Separation of top 5 weighted GBC model features in syn-PSMB2-ψ (GU**ψ**CG, blue) syn-PSMB2-U (GU**U**CG, red) synthetic training data. Features are compared with 2D scatter plots (upper-half), histogram distribution for each feature (diagonal), and contour plots (lower-half).

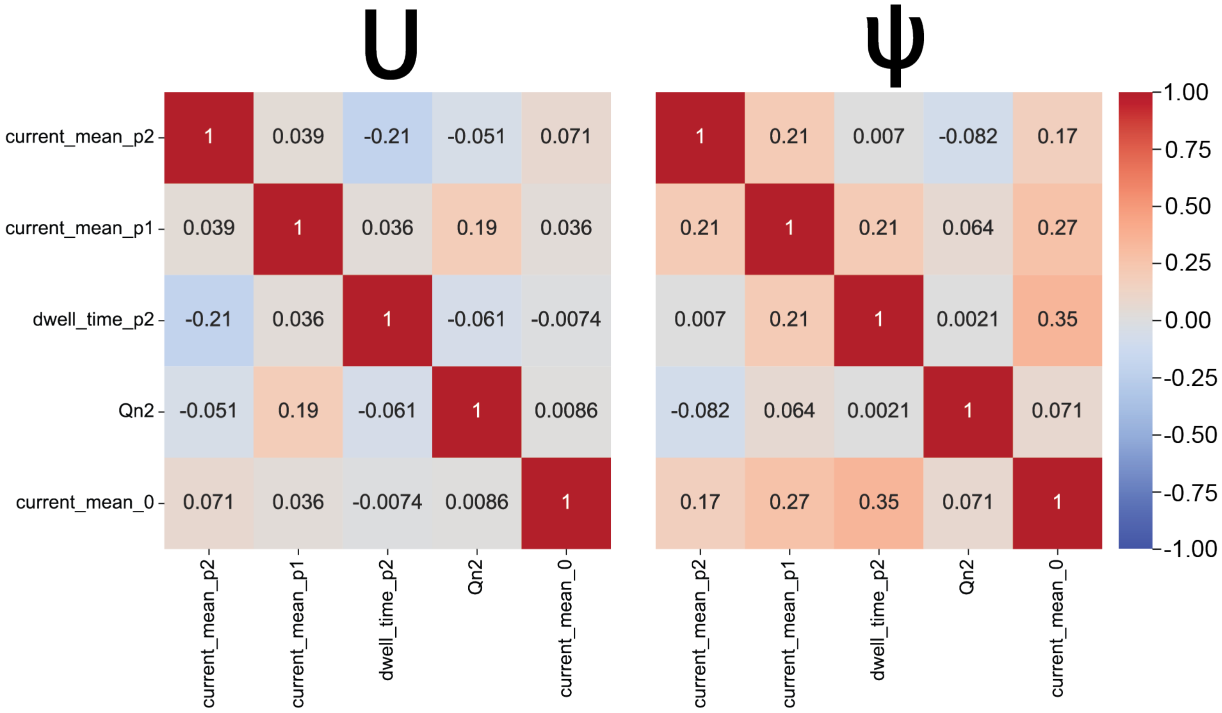

**Figure S8**: Cross-correlation heat map (normalized) of top 5 weighted features in syn-MCM5-U (left) and syn-MCM5-ψ (right) reads after GBC model training and testing.

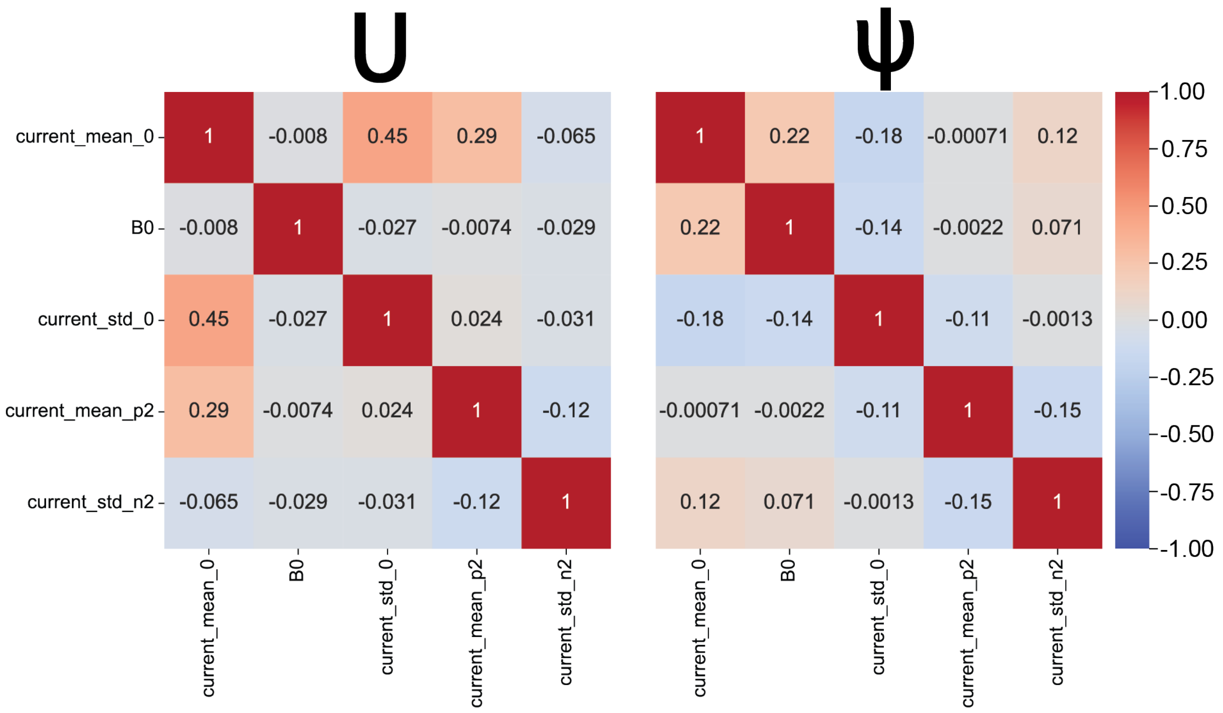

**Figure S9**: Cross-correlation heat map (normalized) of top 5 weighted features in syn-MRPS14-U (left) and syn-MRPS14-ψ (right) reads after GBC model training and testing.

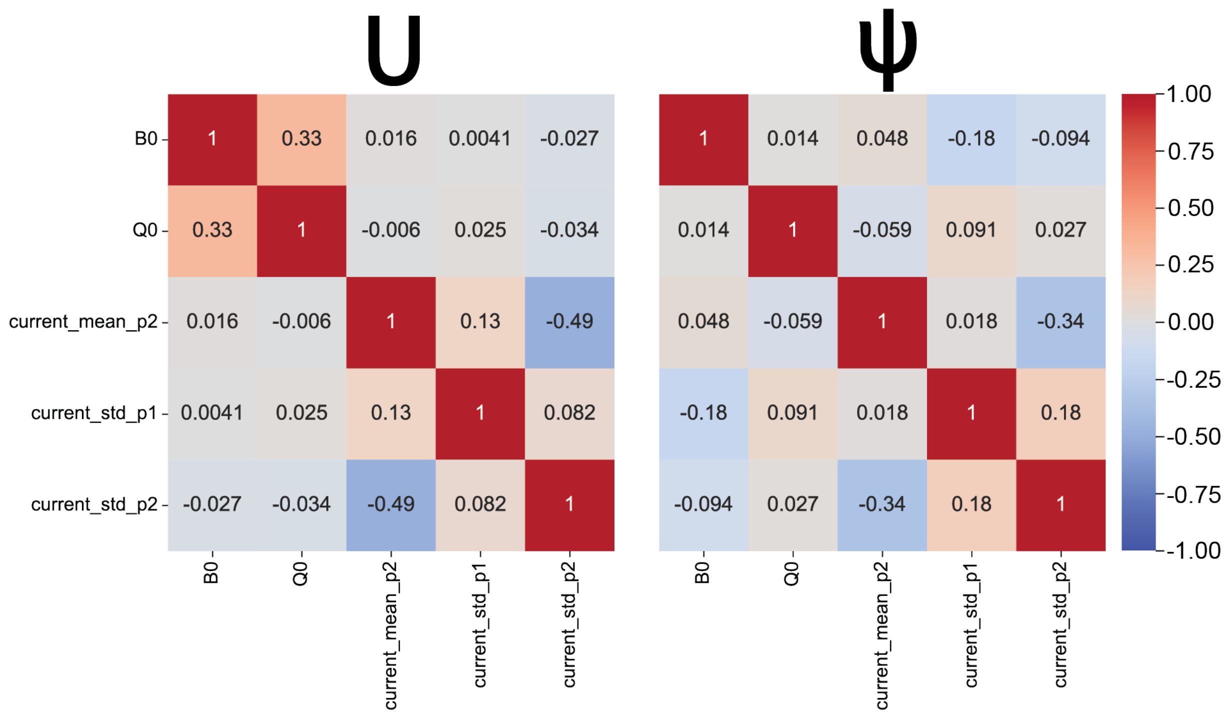

**Figure S10**: Cross-correlation heat map (normalized) of top 5 weighted features in syn-PRPSAP1-U (left) and syn-PRPSAP1-ψ (right) reads after GBC model training and testing.

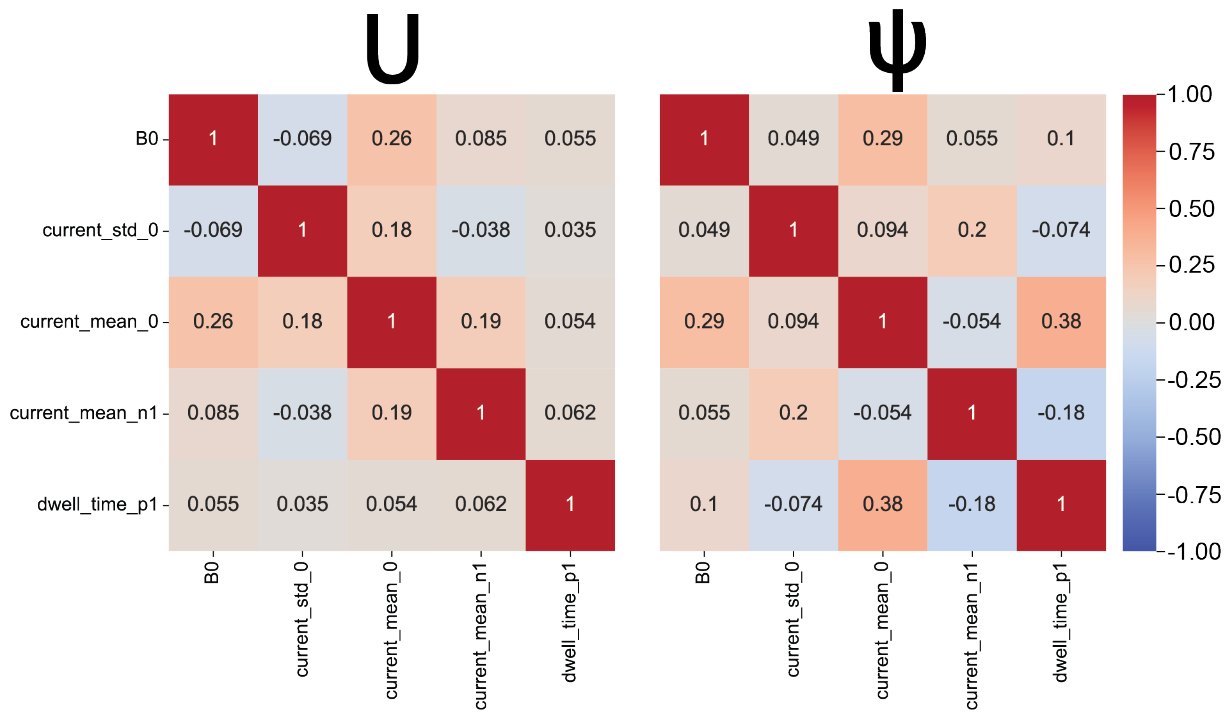

**Figure S11**: Cross-correlation heat map (normalized) of top 5 weighted features in syn-PSMB2-U (left) and syn-PSMB2-ψ (right) reads after GBC model training and testing.

**Supplementary Note 1**: Top 5 weighted features for each synthetic construct (gene) observed during GBC fitting

Except for PRPSAP1, the current mean when ψ is present in k-mer position 0 is a top five weighted feature. One or more current mean of a ψ-containing k-mer is present in all 4 synthetic constructs. K-mer current standard deviation was a top 5 weighted feature for all the synthetic-trained models except for MCM5. Dwell time was only seen in the top five for PSMB2 and MCM5, in particular, the dwell time when ψ is present in k-mer position +1 and +2, respectively. Quality score as a top five feature was only seen in MCM5 (quality score of basecall in position -2 from ψ) and PRPSAP1 (quality score of basecall in the position of ψ). The basecall at the ψ position was in the top five PSMB2, MRPS14, and PRPSAP1. None of the Fourier components were present in the top 5 weighted features for all 4 synthetic constructs.
